# MocFormer: A Two-Stage Pre-training-Driven Transformer for Drug-Target Interactions Prediction

**DOI:** 10.1101/2023.09.13.557595

**Authors:** Yilun Zhang, Wentao Wang, Jiahui Guan, Deepak Kumar Jain, Tianyang Wang, Swalpa Kumar Roy

## Abstract

Drug-target interactions (DTIs) is essential for advancing pharmaceuticals. Traditional drug-target interaction studies rely on labor-intensive laboratory techniques. Still, recent advancements in computing power have elevated the importance of deep learning methods, offering faster, more precise, and cost-effective screening and prediction. Nonetheless, general deep learning methods often yield low-confidence results due to the complex nature of drugs and proteins, bias, limited labeled data, and feature extraction challenges. To address these challenges, a novel two-stage pre-trained framework is proposed for DTIs prediction. In the first stage, pre-trained molecule and protein models develop a comprehensive feature representation, enhancing the framework’s ability to handle drug and protein diversity. This also reduces bias, improving prediction accuracy. In the second stage, a transformer with bilinear pooling and a fully connected layer (FCN) enables predictions based on feature vectors. Comprehensive experiments were conducted using DrugBank dataset and Epigenetic-regulators dataset to evaluate the framework’s effectiveness. The results demonstrate that the proposed framework outperforms the state-of-the-art methods regarding accuracy, area under the ROC curve (AUC), recall, and the area under the precision-recall curve (AUPRC). The code will be available after being accepted: https://github.com/rickwang28574/MocFormer

## 1. Introduction

Predicting the drug-target interactions (DTIs) could be applied in multiple fields, for example, the drug discovery [1], drug repositioning [2], and the prediction of drug side effect [3]. The critical drug discovery process is identifying DTIs among numerous candidates[4]. Although conventional measurement in vitro experimental testing can verify DTIs, it suffers from extremely long time and monetary costs. To reduce the wet-lab-based verification procedure’s expensive workload, computational approaches are adopted to efficiently filter potential DTIs from a large number of candidates for subsequent biological experiments[5]. The traditional in-silico computational methods could generally be classified into three categories: ligand-based, target-based, and chemogenomic approches [6]. The ligand-based approaches predict the DTIs by the similarities of the ligands. They search for similar compounds verified to interact with the particular target. Target-based methods claim that a target with a similar 3D structure can interact with the same drug. However, both methods rely on powerful computing resources, running time, and accurate 3D protein structures. With the development of machine learning and deep learning in recent years, chemogenomic methods have begun to be widely used. These use target and ligand characters simultaneously to make the interaction predictions. These computational approaches rely on machine learning techniques to build a prediction model to accurately estimate undiscovered interactions based on the chemogenomic space that incorporates drug and target information.

Recently, various deep models have shown encouraging performance in DTI predictions. Initially, researchers usually only used manual annotation to label proteins and small molecules with manual descriptors in limited datasets. Then, researchers proposed a CNN-based model [7], which utilizes multi-scale one-dimensional convolutional neural network [8] to obtain targets features and Extended Connectivity Fingerprints [9] to get compounds features. Also, the attention mechanism is introduced into DTIs prediction [10,11]. At the same time, with the further development of deep learning [12–17], transformer [18] and GNN [19] were proposed, and attempts were made to encode and decode molecules and proteins separately through transformer [20]. Encoding and decoding [20] to learn their high-dimensional structures and input them into neural networks for iteration to simulate their interactions. Meanwhile, graph neural networks are also the usual means to study DTIs, where one constructs its 2D structure by treating atoms as nodes and chemical bonds as edges. The attention mechanism has been widely used in both approaches, which is thought to capture the key sites where its small molecules bind to proteins [21]. Hyper-AttentionDTI’s attention mechanism can infer the interactions of each amino acid atom pair but also control the characteristics on the channel [22]. DrugBAN proposed a bilinear attention network with domain adaptation to explicitly learn pairwise local interactions between drugs and targets and has a specific generalization ability. Both methods can represent local interactions to some extent through improved attention mechanisms. In recent years, with the development of molecule and protein language pre-training models, people have tried to encode the smiles and protein sequences of molecules into vectors to represent their physical and chemical functions and structural information [23–27]. These pre-trained models are trained on an extensive unlabeled molecule and protein data set so these vectors can better represent their physical and chemical characteristics. DeepLPI [28] and AI-Bind [29] utilize these pre-trained models to make the DTI predictions.

Despite these efforts, the following challenges are still open. 1) The complex nature of drugs and proteins presents a formidable challenge. These molecules exhibit various structural variations, chemical interactions, and biological functions, making them difficult to predict accurately. 2) Inherent bias in the data can introduce significant uncertainties. Biomedical datasets are often collected from specific populations or experimental conditions, which may not fully represent the diversity of biological systems. This bias can result in models that perform well in particular scenarios but struggle when applied to more diverse or real-world situations. 3) A limited availability of labeled data poses a substantial challenge. Supervised learning methods rely on labeled examples for training and require substantial quantities of accurately annotated data. In drug discovery and protein analysis, obtaining large, high-quality labeled datasets is expensive and time-consuming. Consequently, models may not be sufficiently trained to handle the full spectrum of potential inputs, leading to lower confidence in their predictions. 4) Feature extraction remains a persistent challenge. Identifying and selecting relevant features from complex biological data is a non-trivial task. Inaccurate feature representation or the omission of crucial information can significantly impact the performance of predictive models, contributing to the uncertainty in their results.

To overcome the above issues, a two-stage framework is proposed for accurate and rubust drug-target interactions (DTIs) prediction, as shown in Fig. 1. In the first stage, pre-trained molecule and protein foundation models are applied to encode the drug and protein sequences into comprehensive feature vectors. The advantages of the pre-training are twofold. Firstly, pre-training foundation models for molecule and protein structures provide a powerful starting point for feature representation. This pre-trained model addresses the limitation of having a limited number of labeled protein-drug pairs when using deep learning methods to predict drug-target interactions (DTIs). Compared to previous deep learning methods which builds embeddings from a limited number of proteins and molecules using relatively simple methods from scratch, the approach of using transfer learning with pre-trained models for molecule and protein representation allows training on a much larger dataset of unlabeled molecules and proteins, thereby avoiding overfitting and obtaining more accurate features. In the second stage, a transformer with bilinear pooling and a fully connected layer further processes the feature vector acquired from the first stage and outputs the final prediction result. It enhances the grasp of drug-target relationships and interaction prediction accuracy, spanning various scales like molecule structures.

**Figure 1.**
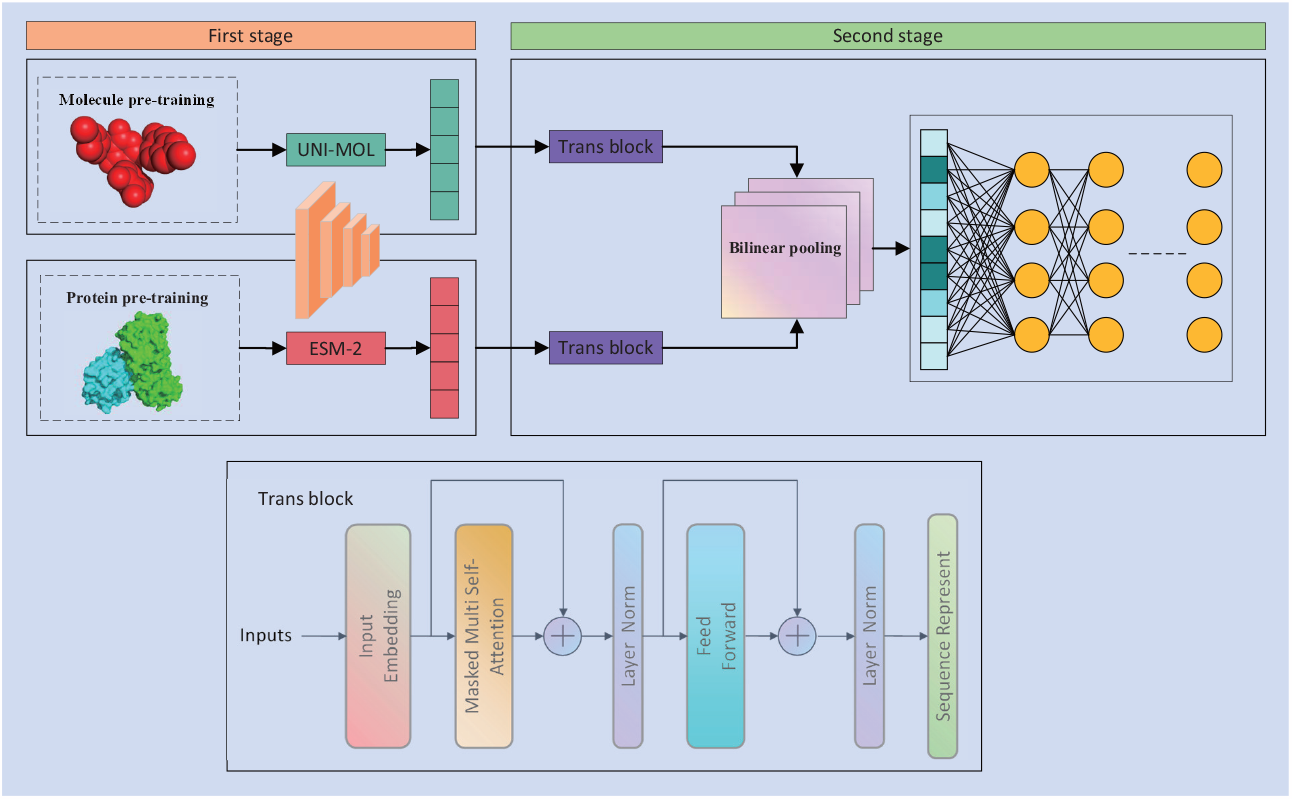
The overarching workflow of the proposed framework encompasses three pivotal constituents: a data representation driven by pre-training, a transformer influenced by pre-trained models, and the dissemination of results.

In summary, this paper presents the following contributions: 1) To the best of our knowledge, a pre-training driven transformer framework is proposed for the first time, termed MocFormer, to achieve drug and target interactions prediction based on transfer learning. 2) The first stage obtains a comprehensive vector representation of molecule and protein features through fine-tuning and transfer learning. The second stage enhances the grasp of drug-target relationships and the accuracy of interaction prediction through transformer, bilinear pololing and FCN. 3) Experimental results show that our method outperforms the most recent state-of-the-art drug-target interactions methods on two public benchmarks, demonstrating the effectiveness of our method and its potential applicability in clinical practice.

The structure of this paper is outlined as follows: In Section 2, we provide an overview of the framework by presenting its workflow. Section 3 presents the experimental setup and comprehensive experimental results. In Section 4, we summarize the entire paper and give priorities for future work.

## 2. Methods

Figure 1 offers a comprehensive illustration of the framework for identifying drug–target interactions (DTIs) through the utilization of drug SMILES strings and protein amino acid sequences. The framework is divided into two primary stages. In the first stage, we employ pre-trained foundational models to process molecule and protein data. In the subsequent second stage, we utilize a pre-training-driven transformer, bilinear pooling, and a fully connected layer (FCN) to further refine the DTIs prediction. This two-stage process is pivotal in achieving accurate and reliable predictions of drug–target interactions.

### 2.1. Molecule Pre-trained Module

Uni-Mol is an advanced framework designed for 3D molecule representation learning with three key components. 1) The foundation of Uni-Mol relies on a transformer-based backbone. This backbone effectively processes input data consisting of individual atoms and atom pairs, integrating the SE(3) methodology to condense the intricate 3D structure of molecules effectively; 2) To ensure robustness and comprehensive learning, Uni-Mol undergoes training on a vast dataset, encompassing an impressive 209 million molecules and 3 million proteins. This extensive training dataset equips the model with a broad understanding of molecule structures and their relationships; 3) Uni-Mol’s capabilities are further enhanced through fine-tuning various downstream tasks. These tasks include predicting drug-target interaction sites, distinguishing between correct and incorrect binding sites, and predicting the corresponding 3D structures. Fine-tuning refines the model’s abilities to make precise predictions and contributes to its overall versatility in molecule analysis.

In the MocFormer pipeline, the grid search method was employed to fine-tune the pretrained model provided by Uni-Mol on the DrugBank dataset. The pre-trained model from the DrugBank dataset underwent fine-tuning using the random forest regression method, and the learning rate was selected from the range [1e-5, 1e-4, 4e-4, 1e-3]. Furthermore, different batch sizes, namely [8, 16, 32], were experimented with. To ensure robustness, the five-fold cross-validation technique was utilized. This technique allowed for the selection of three sets of optimal characterization results. These optimal sets of representation vectors were then used as input for MocFormer’s model inference, and the final choice was determined based on the best performance. Following the preprocess of the Molecule Pre-trained Module, we obtain the drug’s embedding matrix, which is denoted as *f*_*D*_. This computation can be concisely expressed using Equation (1), where *f* denotes the size of the embeddings for drug strings, and we’ve set this dimension to 512. This equation encapsulates the transformation of drug data into a meaningful embedded representation. The result of molecule pre-training is illustrated in Figure 2, which presents a word cloud visualization.

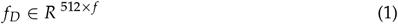

**Figure 2.**
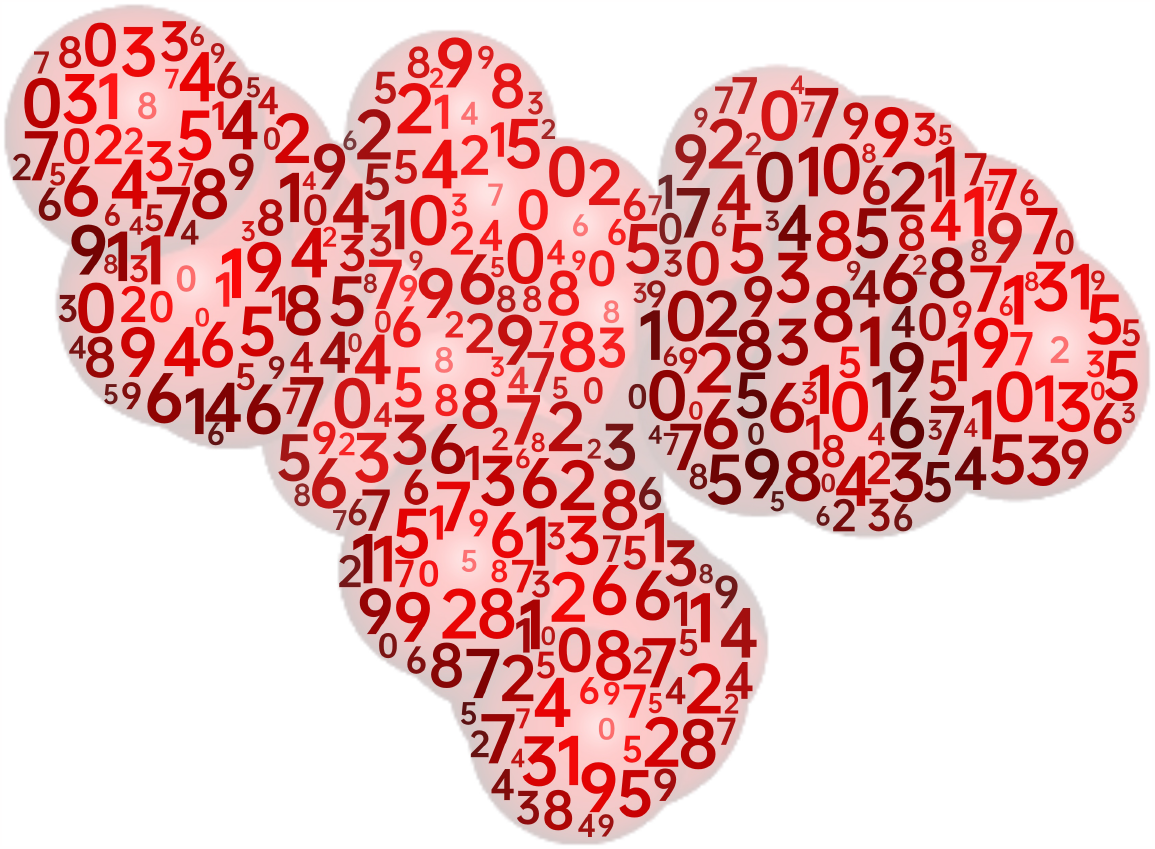
Word cloud visualization of molecule pre-training results.

### 2.2. Protein Pre-trained Module

ESM-2 is developed based on the belief that the information regarding structure and function can be found in amino acid sequences, making LLMs (Large Language Models) a handy tool for this task. ESM-2 remains a transformer-based model with a maximum of 15 billion parameters. It utilizes approximately 138 million sequences for training and employs an equivalent transformer to represent the protein’s three-dimensional structure. This results in an attention pattern corresponding to the protein’s three-dimensional structure.

In the MocFormer model, the chosen variant of ESM-2 is a large language model with 36 layers and 3 billion parameters. A fine-tuning process is meticulously designed to enhance its capabilities further and adapt it for the Drug-Target Interactions (DTIs) task. The selected method for fine-tuning is the K-neighborhood algorithm, which is optimized using the grid search approach. The hyperparameters being searched include the batch size (options: 8, 16, 32), the number of neighbors (options: 5, 10), the weighting strategy (options: uniform, distance), and the algorithm type (options: ball_tree, kd_tree, brute). The leaf size is also considered for the algorithm (options: BallTree, KDTree). Finally, three distinct sets of vector representations are derived. These sets are then utilized as input for the subsequent model in the pipeline. The goal is to identify the set of representation vectors that consistently delivers the best performance, ensuring that the final model is optimized for the DTIs task. After the Protein Pre-trained Module processes the input, the protein’s embedding matrix, denoted as *f*_*P*_, is obtained. The computation can be summarized using Equation (2), where *f* represents the size of the embeddings for protein strings, and 2560 means the embedding dimensions. The result of protein pre-training is illustrated in Figure 3, which presents a word cloud visualization.

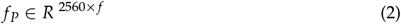

**Figure 3.**
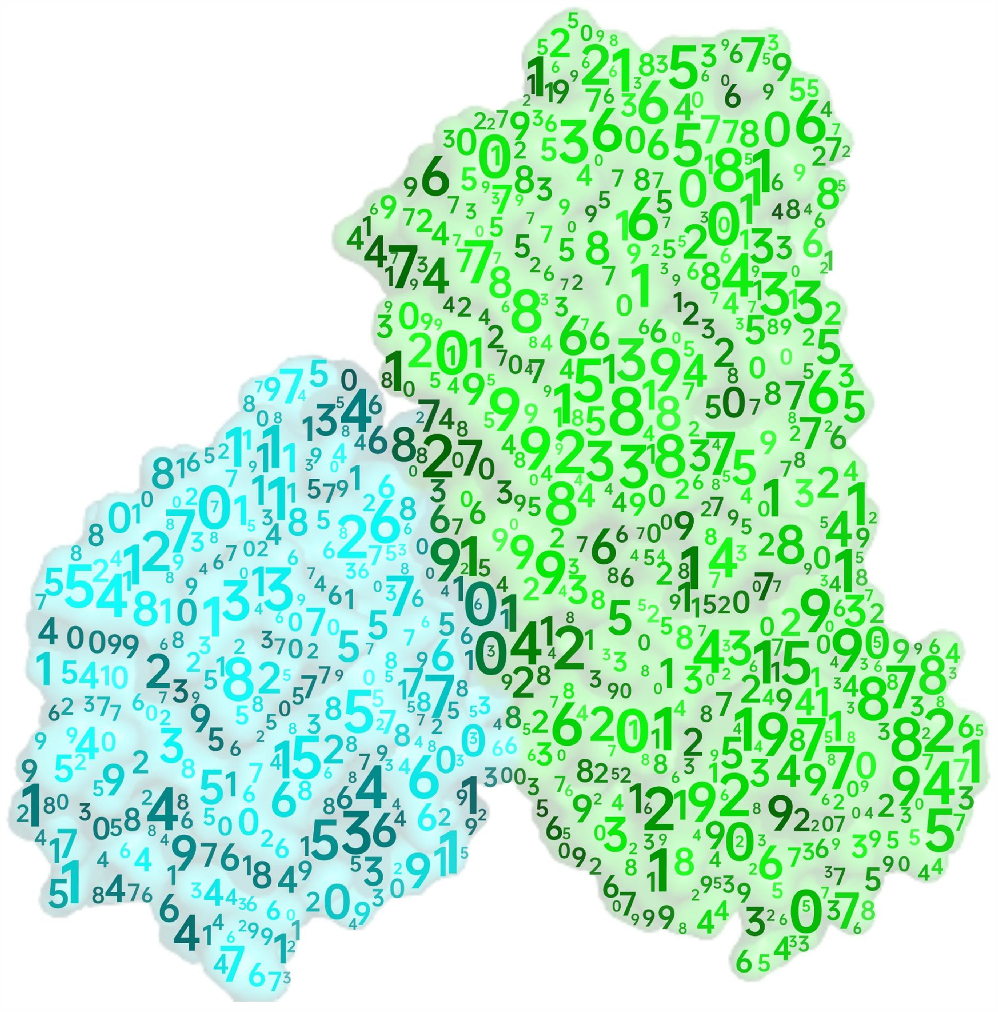
Word cloud visualization of protein pre-training results.

### 2.3. Transformer Module

In this pipeline, transformer modules utilize a multi-attention mechanism to calculate the feature vector of molecule and protein acquired from the first stage. This mechanism assigns weights to dimensions from the 512-dimensional drug vectors and the 2,560-dimensional protein vectors. Moreover, the multi-head attention mechanism within the transformer further improves this process, ensuring that the more critical vector dimensions are focused on. The allocation of weights facilitates MocFormer in learning the intrinsic patterns associated with drug-target interactions. By focusing on the most pertinent vector dimensions, MocFormer learns and captures the intrinsic relationships and nuances inherent in drug-target interactions.

The computation can be summarized using Equation 3–6. Where *Q* represents the query, *K* represents the key, and *V* represents the value. The weight matrices are denoted as *W*^*Q*^, *W*^*K*^, and *W*^*V*^, while *d*_*k*_ means the dimensions of the vectors.

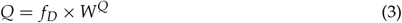

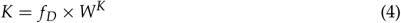

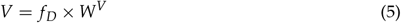

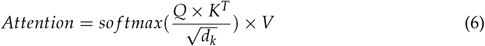

The multi-head attention is then introduced and summarized using Equation 7–8. For each head, there are weight matrices 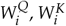, and 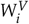, with dimensions 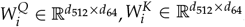, and 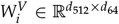. Additionally, a linear transformation matrix 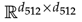 is utilized.

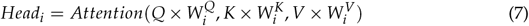

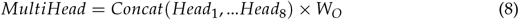

The fully connected feed-forward network comprises two dense layers, each followed by a ReLU activation function, allowing for nonlinear transformations. This can be summarized using Equation 9. The weight matrices *W*_1_ and *W*_2_ have dimensions of ℝ^*f × f*^, and bias terms *b*_1_ and *b*_2_ are also included.

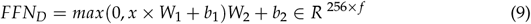

The similar process is used for protein embedding vectors.

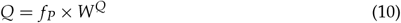

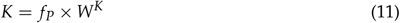

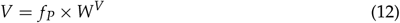

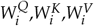 are all weight matrices, 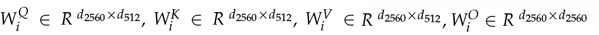 is the linear transformation matrix.

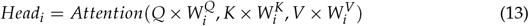

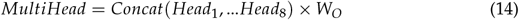

Also, the network comprises two dense layers, each followed by a ReLU activation function, allowing for nonlinear transformations. This can be summarized using Equation 15. The weight matrices *W*_1_ and *W*_2_ have dimensions of ℝ ^*f × f*^, and bias terms *b*_1_ and *b*_2_ are also included.

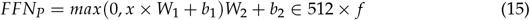

### 2.4. Bilinear Pooling and Full Connected Layer

The bilinear pooling technique fuses features from the drug and protein decoders. It involves bilinearly multiplying the first two features at the same position to obtain the matrix **B**. Then, sum pooling is applied to all positions in **B** to get the matrix *ξ*. The matrix *ξ* is further transformed into a vector, referred to as the bilinear vector **x**. Additionally, moment normalization and L2 normalization operations are performed on **x** to obtain the fused features **Z**. The bilinear pooling method is utilized to merge the output of the drug and protein decoders. Then, the merged vector representation will be fed into a multi-layer, fully connected layer network. The activation function is relu, a dropout layer is added after each layer to prevent overfitting, and a binary cross entropy is used to output the final prediction results. The specific calculation process can be expressed by Equation 16–20.

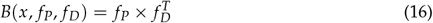

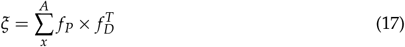

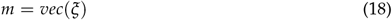

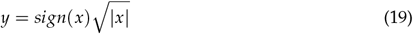

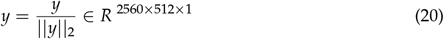

## 3. Experimental Results

This section presents the results obtained by applying the proposed methods to the DrugBank dataset. The experimental dataset and evaluation metrics will be explained in Section 3.1. The implementation details of the experiments will be discussed in Section 3.2. In addition, Section 3.3 will present the results of the ablation study, while Section 3.4 will provide a comprehensive comparison with the current state of the art.

### 3.1. Dataset and Evaluation Metrics

#### DrugBank dataset

The experimental dataset for this study was derived by extracting drug and target data from the DrugBank database [30], as presented in Table 1. The dataset used in this research corresponds to the data released on January 3, 2020 (version 5.1.5). Inorganic compounds and tiny molecule compounds (e.g., Iron [DB01592] and Zinc [DB01593]) were manually discarded, along with drugs having SMILES strings that could not be recognized by the RDKit Python package [31]. After this filtering process, 6,655 drugs, 4,294 proteins, and 17,511 positive drug-target interactions (DTIs) remained in the dataset. To create a balanced dataset with equal positive and negative samples, unlabeled drug-protein pairs were sampled following a common practice [20,32]. This approach allowed for the generation of negative samples, resulting in a balanced dataset for analysis.

**Table 1.** Summary of the DrugBank dataset.

| Datasets | Protein | Drug | Interaction | Positive | Negative |
| --- | --- | --- | --- | --- | --- |
| DrugBank | 4294 | 6655 | 35022 | 17511 | 17511 |

#### Epigenetic-regulators dataset

This dataset is based on protein family-specific datasets (Large-scale) [33], further constructed by applying a strategy that only considers compound similarities while distributing bioactivity data points into train-test splits, as presented in Table 2. Compounds in train and test splits are dissimilar (Tanimoto score *<* 0.5). Therefore, similar compounds cannot participate in both train and test splits. This strategy makes the prediction task more challenging and realistic than random splitting. It partly prevents the model from memorizing bioactivities over identical or highly similar compound fingerprints shared between train and test folds. In previous experiments of a similar nature, researchers segregated distinct molecules into the training and test sets. In other words, they ensured that the molecules in the test set were not present in the training set and vice versa. However, in our current study, we have taken a different approach by including entirely dissimilar types of molecules. Specifically, molecule A is part of the test set, while molecule B is found in the training set. These molecules exhibit a substantial dissimilarity, indicated by a Tanimoto score of less than 0.5. This implies that their degree of similarity is exceedingly low, akin to the difference between humans and dogs, as opposed to mere similarity, such as that between men and women. The score quantifies the degree of similarity. The term “Pchem value” denotes the experimental measurement of the interaction between the target and ligand. In this context, we selected a threshold of 6. If the Pchem value exceeds 6, it indicates the presence of an interaction, and the corresponding label is set to 1.

**Table 2.** Summary of the epigenetic-regulators dataset.

| Datasets | Protein | Drug | Interaction | Positive | Negative |
| --- | --- | --- | --- | --- | --- |
| Epigenetic-regulators | 113 | 10120 | 17757 | 8834 | 8923 |

Four key metrics were considered for a comprehensive performance analysis: Accuracy, AUC, Recall, and Area Under the Precision-Recall Curve (AUPRC). Accuracy assesses overall correctness, AUC evaluates the model’s ability to rank positive and negative samples correctly, Recall measures the model’s effectiveness in identifying positive samples, and AUPRC evaluates the model’s performance in classifying imbalanced datasets.

### 3.2. Implementation Details

The framework used in this study is built on the PyTorch platform and utilizes an NVIDIA Tesla V100S GPU. The entire dataset was divided into training, validation, and testing sets, with proportions of 70%, 20%, and 10%, respectively. Each experiment employed a 5-fold cross-validation approach. The AdamW optimizer optimized the model with an initial learning rate of 0.001 and a weight decay 0.001. Additionally, a learning rate schedule based on ReduceROnPlateau was implemented. This schedule had a patience of 5, meaning that if the model’s validation loss did not decrease after five epochs, the learning rate would decay to 10% of the previous rate.

### 3.3. Ablation Study

To assess the effectiveness of each component in our method, a series of ablation experiments were conducted, as presented in Table 3 and Figure 4. These experiments progressively enhanced the baseline network by applying the following configurations: 1) Adding only the molecule pre-trained module (A) to the baseline. 2) Adding only the protein pre-trained module (B) to the baseline. 3) Simultaneously adding the molecule and protein pre-trained modules to the baseline. 4) A transformer with bilinear pooling (C) was incorporated after combining the molecule and protein pre-trained modules with the baseline. **Baseline**: The baseline will be to perform the encoding and decoding process for small molecules and protein sequences using the two word2vec functions in gensim, respectively, using average pooling to connect the two types of vectors and pass them to the multilayer perceptron MLP (consisting of multiple fully-connected layers). **Baseline+A**: Baseline+A will replace the original word2vec representation for molecule in baseline with the pre-trained model (fine-tuned) of Uni_mol to characterize small molecules vectorially. At the same time, proteins are still processed using word2vec. Other settings are the same as the baseline. **Baseline+B**: Baseline+B will replace the original word2vec representation for protein in baseline with the pre-trained model (fine-tuned) of ESM-2 to generate embeddings. At the same time, molecules are still processed using word2vec. Other settings are the same as the baseline. **Baseline+A+B**: Although Uni_mol and ESM-2 are known as powerful molecule characterization models, using word2vec-generated vector representations as input might cause the model to rely on topology for predictions, thereby lacking practical biochemical meaning. This issue influences interactions between the single-side (Molecule or Protein) word2vec module and the pre-trained molecule characterization module (Protein or Molecule), causing them to learn an incorrect paradigm and ultimately resulting in weakened results. Therefore, better performance can be achieved by simultaneously pre-training encoding for both molecules and proteins. **Baseline+A+B+C**: Our final framework is Baseline+A+B+C. In this framework, Uni_mol and ESM-2 generate the embeddings, which are then input into the MLP after passing through the transformer and bilinear pooling layers, ultimately yielding the prediction results.

**Table 3.**
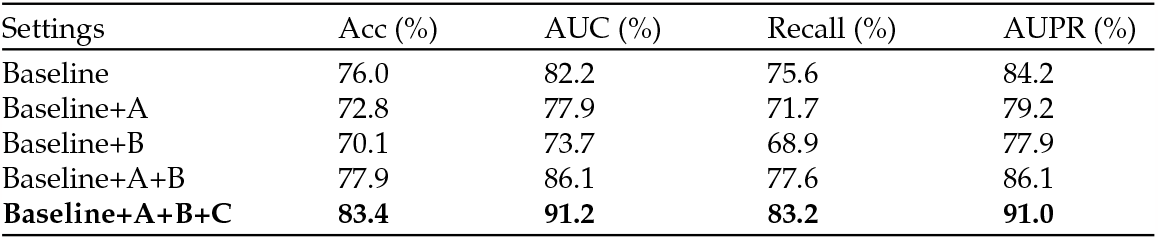
Results of ablation studies.

**Figure 4.**
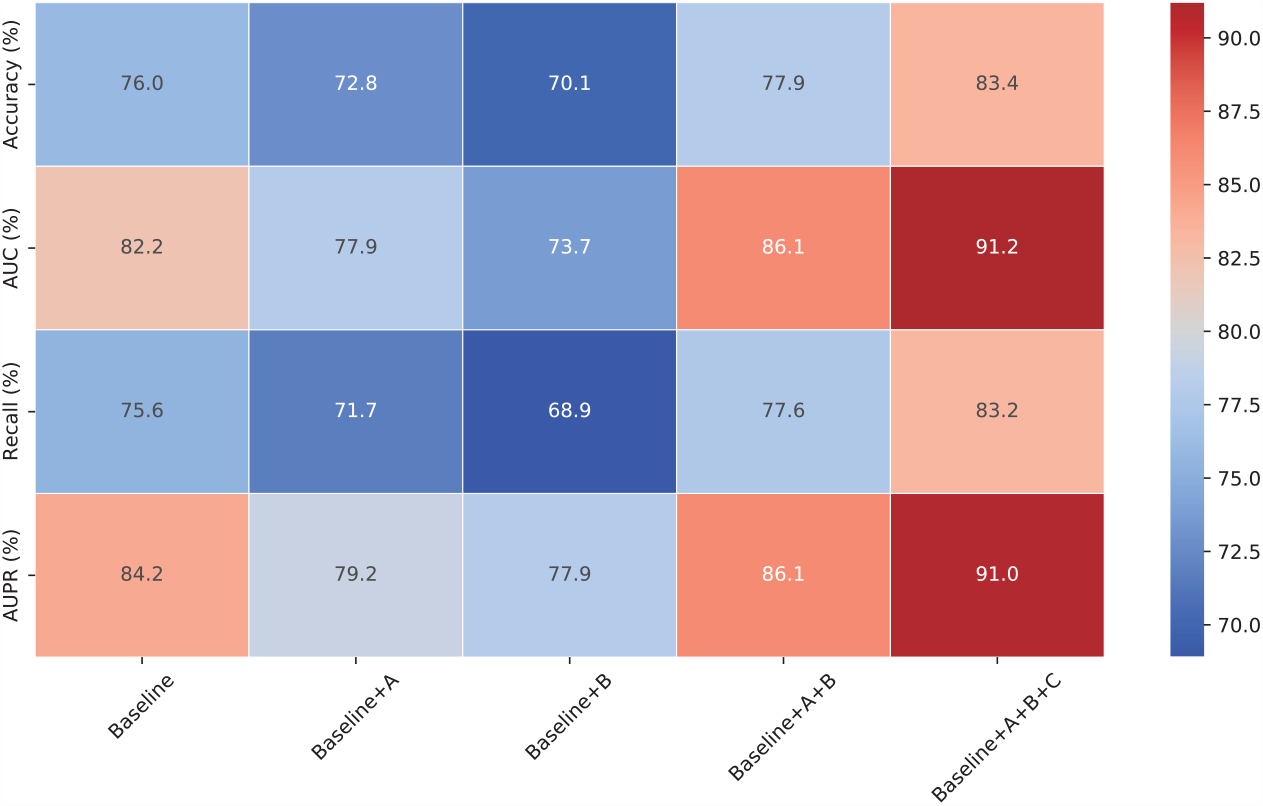
The heatmap shows that adding conditions “A” and “B” individually initially reduces performance, but combining them yields a significant positive impact (“Baseline+A+B”). Moreover, introducing condition “C” ultimately allows the model to achieve the best performance.

### 3.4. Comparison with the State-of-the-Art

To establish the superiority of our proposed method, we conducted comprehensive comparison experiments, pitting it against two attention-based networks (DrugBAN and HyperAttentionDTI), one transformer-based network (Moltrans), and one transfer-learning-based network (AI-Bind). These experiments were carried out using the DrugBank dataset. The results demonstrate that our method surpasses previous methods, achieving state-of-the-art (SOTA) performance across multiple critical evaluation metrics, including accuracy, AUC, recall, and AUPR. These findings are detailed in the data presented within table 4. For further clarity, our method’s dominance across these metrics is visually emphasized in the bar graph displayed in figure 5. This graphical representation vividly illustrates how our approach consistently outperforms other methods in terms of their average scores. Furthermore, the box plot presented in figure 6 underscores our method’s robustness and stability across these four key performance indicators.

**Table 4.**
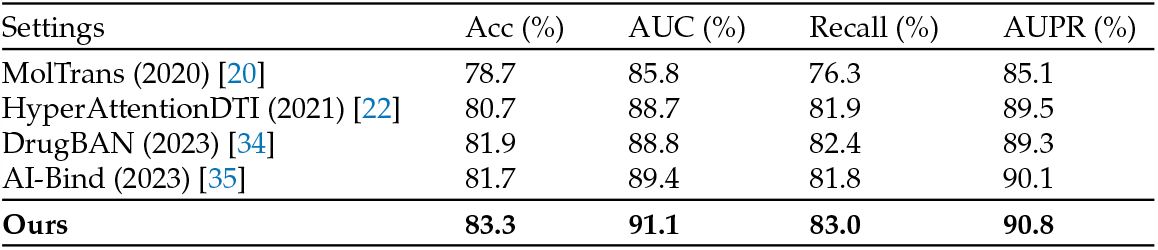
Results of quantitative comparisons on DrugBank dataset.

| Settings | Acc (%) | AUC (%) | Recall (%) | AUPR (%) |
| --- | --- | --- | --- | --- |
| MolTrans (2020) [20] | 78.7 | 85.8 | 76.3 | 85.1 |
| HyperAttentionDTI (2021) [22] | 80.7 | 88.7 | 81.9 | 89.5 |
| DrugBAN (2023) [34] | 81.9 | 88.8 | 82.4 | 89.3 |
| AI-Bind (2023) [35] | 81.7 | 89.4 | 81.8 | 90.1 |
| **Ours** | **83.3** | **91.1** | **83.0** | **90.8** |

**Table 5.**
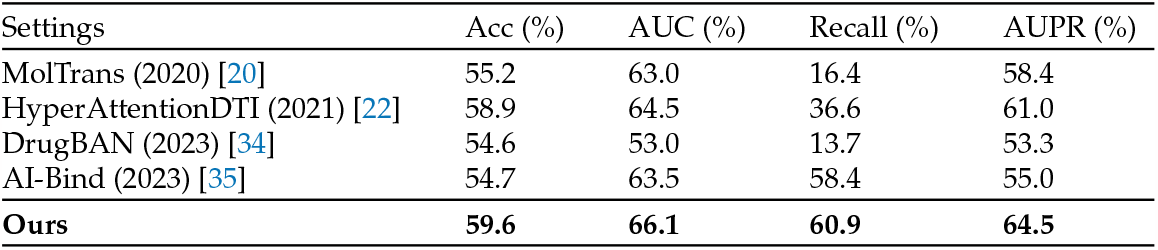
Results of quantitative comparisons on Epigenetic-regulators dataset.

**Figure 5.**
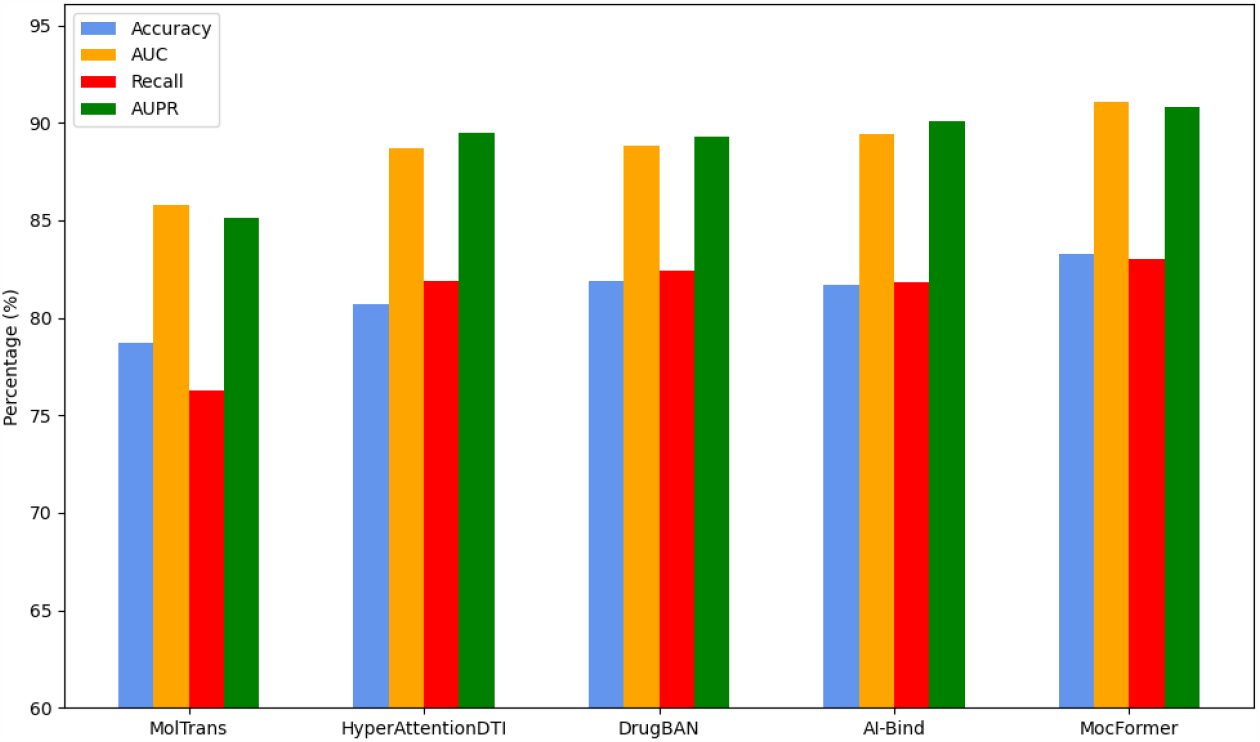
Bar chart visualization of quantitative comparisons on DrugBank dataset.

**Figure 6.**
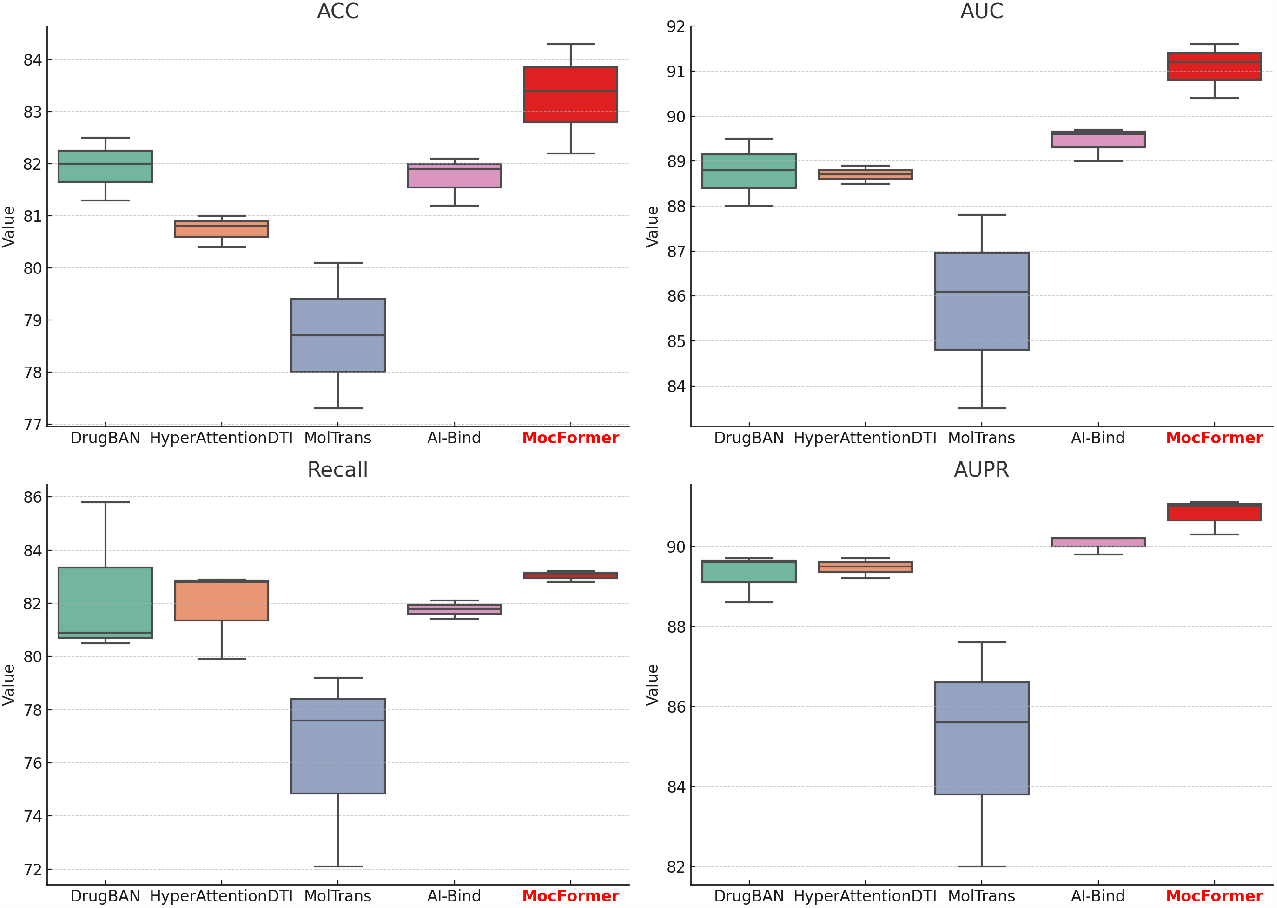
Box chart visualization of quantitative comparisons on DrugBank dataset.

To further showcase the remarkable generalization capabilities of our proposed approach, we conducted supplementary experiments utilizing the Epigenetic-regulators dataset. These additional experiments’ outcomes are presented in tabel 5. The bar chart figure 7, similar to the experiments conducted on the DrugBank dataset, is a compelling testament to our method’s consistently superior performance. The data showcased herein reflects the mean performance of each method across an array of evaluation metrics. Furthermore, the box chart figure 8 underscores the robust nature of our approach. This is evidenced by the minimized fluctuations in the metrics, affirming the stability and reliability of our method. This body of evidence firmly establishes that our method consistently demonstrates formidable predictive capabilities, even when exposed to previously unencountered feature data, when compared against alternative methods.

**Figure 7.**
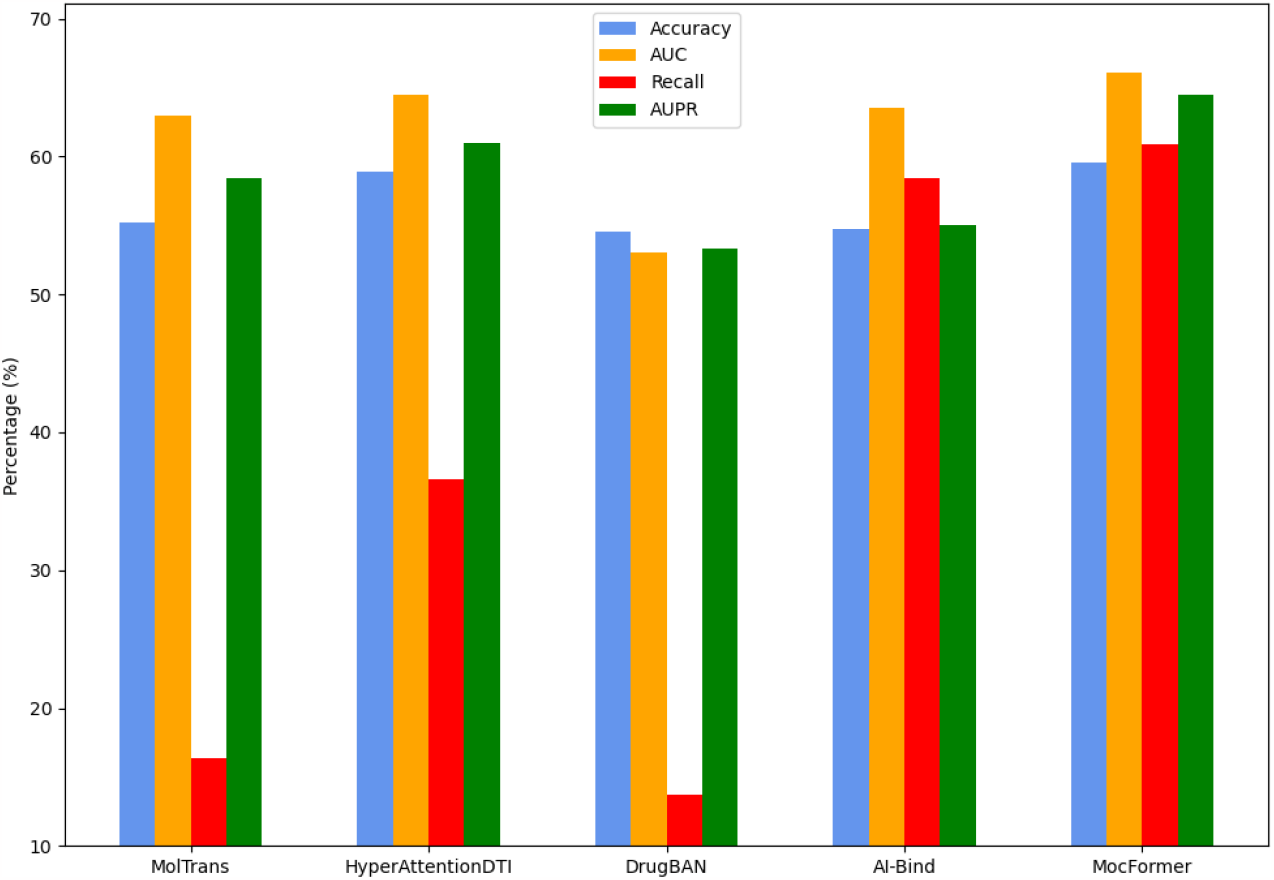
Bar chart visualization of quantitative comparisons on Epigenetic-regulators dataset.

**Figure 8.**
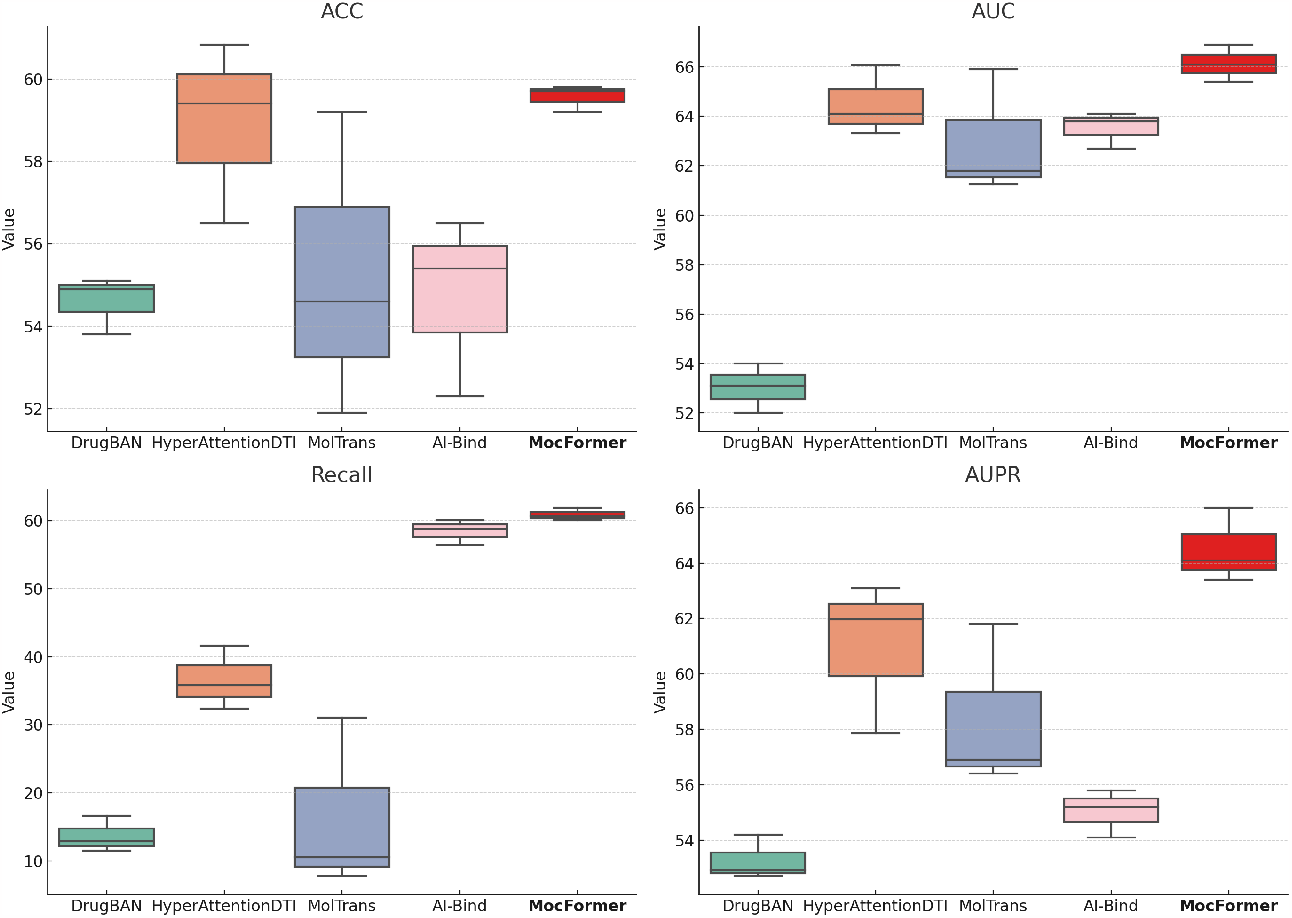
Box chart visualization of quantitative comparisons on Epigenetic-regulators dataset.

## 4. Discussion and Conclusion

This paper introduces a two-stage pre-training driven transformer, a novel framework for identifying Drug-Target Interactions (DTIs). The proposed architecture effectively addresses the challenges posed by the diversity and complexity of drugs and proteins and the presence of bias in the data. Quantitative and qualitative evaluations on the DrugBank and Epigenetic-regulators databases demonstrate that our framework significantly improves accuracy and robustness, achieving state-of-the-art performance. In future work, we would like to investigate the End-to-End Learning Paradigm to optimize the final objective function directly, making it better adapt to the intricacies and complexities of the task process.

## Author Contributions

Conceptualization, Yilun Zhang and Wentao Wang; methodology, Yilun Zhang; software, Yilun Zhang; validation, Yilun Zhang, Wentao Wang and Jiahui Guan; formal analysis, Yilun Zhang; investigation, Yilun Zhang and Jiahui Guan; resources, Yilun Zhang and Jiahui Guan; data curation, Yilun Zhang; writing—original draft preparation, Wentao Wang; writing— review and editing, Wentao Wang, Yilun Zhang, Deepak Kumar Jain, Tian-Yang Wang and Swalpa Kumar Roy; visualization, Yilun Zhang and Wentao Wang; supervision, Yilun Zhang, Wentao Wang, Deepak Kumar Jain, Tian-Yang Wang and Swalpa Kumar Roy; project administration, Yilun Zhang and Wentao Wang; funding acquisition, Yilun Zhang All authors have read and agreed to the published version of the manuscript.

## Institutional Review Board Statement

The study did not require ethical approval for studies not involving humans or animals.

## Informed Consent Statement

Not applicable.

## Data Availability Statement

The two datasets used in the article can be found at [DrugBank](https://github.com/zhaoqichang/HpyerAttentionDTI) and [Epigenetic-regulators](https://github.com/HUBioDataLab/ProtBENCH).

## Acknowledgments

We would like to express our gratitude to all those who helped us during the writing of this manuscript.

## Disclaimer/Publisher’s Note

The statements, opinions and data contained in all publications are solely those of the individual author(s) and contributor(s) and not of MDPI and/or the editor(s). MDPI and/or the editor(s) disclaim responsibility for any injury to people or property resulting from any ideas, methods, instructions or products referred to in the content.

